# The chemopreventive effects of Curcumin against oxidative stress induced by Cadmium or H_2_O_2_ are mediated by Nrf2/ARE signaling and protective autophagy in myeloid cells

**DOI:** 10.1101/2024.07.17.603853

**Authors:** Maria Russo, Annamaria Di Giacomo, Federica Fiore, Carmela Spagnuolo, Virginia Carbone, Paola Minasi, Gian Luigi Russo

## Abstract

The evidence linking high levels of environmental pollutants to chronic degenerative diseases is alarming, with heavy metals (HM) identified as a key factor. Research suggests that certain phytochemicals in the diet can reduce HM levels and mitigate their adverse health effects.Curcumin (Cur), a natural polyphenol, is particularly effective in protecting against Cadmium (Cd) toxicity. The present study demonstrates that preincubation with low doses of Cur (1 μM) in differentiated HL-60 and K-562 human myeloid cells can significantly protect against cytotoxicity induced by Cd and or H_2_O_2_. Cur reduced the increased levels of reactive oxygen species (ROS) generated by Cd or H_2_O_2_ by inducing a protective form of autophagy. Cur activated mild oxidative stress that triggers the expression of Nrf2-dependent transcripts, such as HO and NQO1. The potential chemopreventive effects of Cur against redox stress have been strengthened by the observation that free and unmetabolized Cur is detectable inside the cells after 5 minutes of treatment, and its presence parallels with increased levels of intracellular GSH. These findings suggest that supplementation with Cur in the form of nutraceuticals may represent a promising way to protect people living in highly polluted areas against the adverse effects of HM contaminants.

## 1. Introduction

Heavy metals (HM) such as cadmium (Cd), lead (Pb), arsenic (As), mercury (Hg), and chromium (Cr) are elements that are not biodegradable and are naturally present in various forms throughout the Earth’s crust. Human activities such as agriculture and industry have significantly contributed to environmental issues by causing soil, water, and air pollution. In recent years, the impact of these metals on human health has become increasingly apparent (Rana et al., 2018). Air pollution in the European Union is estimated to cause about 500,000 premature deaths and diseases. The countries with the highest number of deaths attributed to air pollution are Italy, Germany, Poland, France, and the UK (Carvalho, 2019). In Italy, the Campania region’s “Land of Fire” is a stark symbol of a national ecological crisis, significantly tarnishing the region’s image and severely impacting the agrifood sector (Esposito et al., 2018, Triassi et al., 2015). Different studies have shown a concerning increase in heavy metal concentrations, which are high enough to affect the health of animals and humans. This impact includes acute and chronic toxicity, leading to alterations in reproduction, the endocrine system, and immunosuppression (Ebrahimi et al., 2020, Zaccaroni et al., 2014).

Qualified literature emphasizes how various phytochemical compounds, commonly found in the diet, can interact with heavy metals, interfering with their cellular metabolism and aiding in reducing tissue concentrations (Zhai et al., 2015). Curcumin (Cur) is a polyphenol (1,7-bis(4-hydroxy-3-methoxyphenyl)-1,6-heptadiene-3,5-dione) identified in numerous studies as the primary active compound in turmeric. It has been demonstrated to be effective in treating chronic diseases, such as various types of cancer, diabetes, obesity, and cardiovascular, pulmonary, neurological, and autoimmune diseases (Mohajeri et al., 2017, Shanmugam et al., 2015).

Various studies have linked nutritional interventions to protection against health damage caused by HM. For example, in a mouse model of liver toxicity, a combination of 50 mg/kg Cur and 20 mg/kg resveratrol resulted in a reduction of oxidative damage caused by Cd (Eybl et al., 2006). Based on a rat model study, it has been demonstrated that Cur (30 mg/kg) exerted a neuroprotective effect by chelating HM like lead and Cd (Daniel et al., 2004). Curcumin-heavy metal complexes are believed to reduce oxidative stress by lowering reactive oxygen species (ROS) at the cellular and tissue levels (Mohajeri et al., 2017).

Translating animal studies into human clinical trials for nutritional supplements with phytochemicals faces a significant challenge: rapid metabolism and low bioavailability of these compounds after oral ingestion. Although phytochemicals primarily act as antioxidants against ultraviolet radiation-associated cellular damage in plant tissues, various authors hypothesize that dietary phytochemicals, due to their intrinsic chemical instability and low intracellular concentrations, act as “indirect” antioxidants within mammalian cells. Paradoxically, yet realistically, dietary phytochemicals could function as electrophilic molecules inside cells, creating a mild redox imbalance (Shin et al., 2020).

“Moderate” levels of reactive oxygen species (ROS) within cells can function as signaling molecules, altering redox-sensitive amino acids in various proteins. These proteins include phosphatases, kinases, ion channels, and the transcription factor Nrf2. Nrf2 is known as the master regulator of approximately 500 gene transcripts, which play a role in maintaining cellular balance through various interconnected effects. It is believed that Nrf2 enables nucleated cells to adapt and survive in stressful conditions by regulating the expression of different gene networks (Chun et al., 2021). The modulation of intracellular redox status is one of the most well-studied effects. Antioxidant response elements (ARE) are cis-acting DNA sequences located in the region neighbouring phase II antioxidant genes (HO, NQO1). Nrf2 is a transcription factor that binds to ARE, playing a crucial role in the expression of genes mediated by ARE (He et al., 2023). Nrf2 holds high potential for preventing or treating chronic diseases. Several natural phytochemicals, including sulforaphane, Cur, resveratrol, and quercetin, are reported as Nrf2 activators, although the precise mechanisms are yet to be fully understood (Chun et al., 2021, He et al., 2023).

Dietary phytochemicals are considered to possess cancer chemopreventive potential through various mechanisms, often summarized as “pleiotropic action.” One essential process linked to cellular homeostasis in response to internal and external stress is autophagy (“macroautophagy”), which is dedicated to degrading and recycling cellular components. The process involves phagocytosis of damaged macromolecules and organelles by autophagosomes, followed by their digestion in lysosomes, and ultimately, their recycling for efficient cellular metabolism (Russo and Russo, 2018, White et al., 2015). Additionally, autophagy is significantly involved in the pathophysiology of numerous diseases, such as cancer, cardiovascular issues, infectious diseases, and neurodegenerative conditions (Mizushima and Levine, 2020). Autophagy involves several phases regulated by different *ATG* genes, controlling the complete autophagy pathway. The goal is the formation of double-membrane vesicles (phagophore, autophagosome) adorned with LC3 lipidated proteins, known as LC3-II, which subsequently merge with lysosomes. Inside the lysosomes, acidic hydrolases can break down and recycle their cargo (Mizushima, 2020).

A crucial autophagy protein, p62/^SQSTM1^, has multiple domains, including an LIR domain crucial for directing cargo and LC3-II to the autophagosome. Additionally, it can activate Nrf2 and NF-κB through its KIR and TB domains, respectively (Lee et al., 2023, Zhong et al., 2016). The ROS-mediated stress response enhances cell proliferation and induces stress resistance, underscoring the pivotal role of autophagy as a crucial process in stopping cancer formation at its early stage (Russo and Russo, 2018).

Starting from this premise, the accepted hypothesis is that dietary phytochemicals could be “realistic” chemopreventive agents if assumed at the “exact time” and reaching an “effective” intracellular concentration able to counteract environmental stress by activating protective autophagy and/or the Nrf2 antioxidant response (Russo et al., 2017).

The present study aims to verify the efficacy of Cur and explain its mechanism of action in protecting cells against Cd or H_2_O_2_-induced toxicity using nutritional doses (1 μM). It is important to note that Cur is very unstable in an aqueous medium and has very low tissue bioavailability *in vivo* (Shome et al., 2016). We used two *in vitro* models of human myeloid cells obtained after differentiation: i. HL-60 pro-myelocytic blasts treated with vitamin D analog EB1089, inducing myelomonocytic lineage (James et al., 1997); ii. K-562 acute myeloid leukaemia cells treated with resveratrol, inducing erythroid lineage (Della Ragione et al., 2003, Rodrigue et al., 2001).

As far as we know, this study stands out as one of the few to demonstrate the biological activities of Cur at low micromolar concentration (Russo et al., 2024) evidencing that nutritional doses of the molecule can be associated with measurable chemopreventive effects measured by the capacity to establish crosstalk between autophagy and the Nrf2 system in cellular models resembling human peripheral myeloid cells (Tang et al., 2022, Wahyudi et al., 2022).

## 2. Material and Methods

### 2.1 Cell Culture, Differentiation, and Viability Assays

HL-60 and K562 cell lines (EACC) (Gallagher et al., 1979, Lozzio and Lozzio, 1979) were cultured in RPMI medium supplemented with 10% FBS, 1% L-glutamine, and 1% penicillin/streptomycin (EuroClone, Milan, Italy) at 37°C in a humidified atmosphere containing 5% CO_2_. HL-60 cells differentiated into macrophages after one week of supplementing with 100 nM vitamin D (VD) analog EB1082 (Merck/Sigma-Aldrich, Milan, Italy) (James et al., 1997, Miyaura et al., 1985). K-562 cells were differentiated in erythroblasts by adding 50 μM of resveratrol (Merck/Sigma-Aldrich) in the cell culture medium (24-72h) (Della Ragione et al., 2003, Rodrigue et al., 2001). The cells that were already differentiated were pretreated overnight with 1 μM of Cur (Merck/Sigma-Aldrich) or 0.1% dimethyl sulfoxide (DMSO) as a control (Ctrl). The next day, the medium was changed, and the cells were then incubated with 15 μM of Cd Nitrate (Cd (NO_3_)_2_) (Merck/Sigma-Aldrich) or H_2_O_2_ (Merck/Sigma-Aldrich) for an additional 24 h. Cellular viability was assessed using CyQuant dye (Invitrogen, Life Technologies, Milan, Italy) or Trypan Blue (Merck/Sigma-Aldrich) staining as previously described (Russo et al., 2022, Russo et al., 2021). After differentiation, cellular morphology was documented in HL-60 and K-562 by staining with 4-Nitro blue tetrazolium chloride (NBT) (James et al., 1997) () or Neutral Red solution (Merck/Sigma-Aldrich) (Russo et al., 2007). Microphotographs were captured using a fluorescence inverted microscope (Axiovert Zeiss, Milan, Italy) at 400× magnification, utilizing either phase contrast or a fluorescein isothiocyanate (FITC) filter at the end of staining.

### 2.2. Intracellular measurements of ROS and GSH

Cells were incubated at a concentration of 2×10^4^ cells/ml for 30 minutes. During the incubation, two different probes were used: chloromethyl derivative of di-chloro-fluorescein diacetate (CM-DCFDA) at a concentration of 10 µM (Invitrogen, Life Technologies), and dihydroethidium (DHE) at a concentration of 10 µM (Abcam distributed by Prodotti Gianni, Milan, Italy). These probes were used to measure intracellular peroxides and superoxide, respectively. The incubation was carried out in either phosphate-buffered saline (PBS) or wash buffer at 37°C in a humidified 5% CO_2_ atmosphere. The fluorescence was measured using a Synergy HT multiwell reader (Synergy HT BioTek, Milan, Italy) with an excitation of 485 nm and emission of 530 nm for CM-DCFDA, or an excitation of 500-530 nm and emission of 590-620 nm for DHE.

Antimycin A (Merck/Sigma-Aldrich), an inhibitor of complex III of the mitochondrial electron transport chain, was included in the experiment as a positive control for superoxide generation. The results were expressed as DCF fluorescence (%) or DHE fluorescence (%) relative to untreated cells.

To measure intracellular GSH, both untreated and treated cells at a concentration of 4-5×10^4^/ml were placed in PBS with monochlorobimane (50 µM mCB; Merck/Sigma-Aldrich) for 30 minutes. This allowed intracellular GSH S-transferase to create GSH–mCB adducts, which were then detected using fluorescence. After the incubation, the cells were washed, and the fluorescence was measured using a Synergy HT multiwell reader (excitation at 380 nm and emission at 470 nm). The results were expressed as a percentage of mCB fluorescence. Each experimental point was repeated twice in quadruplicate.

### 2.3. Cell Cycle Analysis

HL-60 and K-562 cell lines (1×10^6^/ml) were induced to differentiate by treatments with the Vitamin D analog, EB1089, for 168 (HL-60) or resveratrol for 24 h. First, cells were harvested and counted using a 0.4% Trypan blue solution. Then, the cells were washed in PBS and underwent ice-cold 70% ethanol fixation. After two additional washes in cold PBS, the cells were stained using 50 µg/ml propidium iodide in the presence of 100 µg/ml DNase-free RNAase A for 1 h at 37°C in the dark. Stained cells were analyzed using a flow cytometer (BD FACSCelestaTM, Becton Dickinson, USA), and the FlowJo V10.8.1 software was used to calculate the proportion of diploid cell distribution across different cell cycle phases (Iacomino et al., 2001).

### 2.4 Autophagy Assay

Autophagy activation was assessed using the Cyto-ID autophagy detection kit (Enzo Life Sciences, Milan, Italy), which utilizes a cationic amphiphilic tracer to accurately count the number of autophagosomes inside the cells. The 488 nm-excitable green dye is designed to minimize staining of lysosomes while showing bright fluorescence when incorporated into pre-autophagosomes, autophagosomes, and autolysosomes. After incubation, the cells were washed, and then a mixture of the autophagy detection marker (Cyto-ID) and nuclear dye (Hoechst 33342) was added. The cells were washed with assay buffer and then observed under a fluorescence microscope (Zeiss Axiovert 200; 400× magnification). The number of autophagosomes was quantified by comparing the green fluorescence (Cyto-ID) to the blue fluorescence (Hoechst) using a microplate fluorescence reader (Synergy HT BioTek).

### 2.5. Immunoblotting

The cells were lysed using a lysis buffer that contained protease and phosphatase inhibitors, as reported by (Russo et al., 2022). After measuring the protein concentration using the Lowry method (Peterson, 1979), protein lysates (20 µg) were mixed with 2X Laemmli loading buffer. The sample was heated at 95°C for 5 minutes before being applied to a 7.5, 10, or 12% SDS-PAGE at pH 8, following the established protocol (Russo et al., 2023). The immunoblots were conducted using Polyvinylidene Fluoride (PVDF) membranes and were incubated for around 16 h with primary antibodies such as anti-p27^KIP1^ (Cell Signaling; distributed by EuroClone; cod. #3698), anti cdc2 (Santa Cruz Biotechnologies, Heidelberg, Germany; code # sc-53), anti-Lamin B (Santa Cruz Biotechnologies; code #sc-374015), (anti-p62^SQSTM1^Cell Signaling; code #5114S), anti-NQO1 (Cell Signaling; code #3187) anti-Nrf2 (Cell Signaling; code #12721) and anti-α-Tubulin (Merck/Sigma-Aldrich; code #T9026). The primary antibodies were diluted 1:1000 in Tween20-Tris Buffer Saline (T-TBS) containing 3% Bovine Serum Albumin (BSA). After washing the membranes in T-TBS, they were incubated for 2 h with a horseradish peroxidase-linked secondary antibody raised against mouse or rabbit. The secondary antibody was diluted to 1:20,000 in T-TBS. The immunoblots were developed using the ECL Prime Western blotting detection system kit from GE Healthcare (Milan, Italy). The band intensities were quantified and expressed as optical density using a Gel Doc 2000 Apparatus from Bio-Rad Laboratories (Milan, Italy) and MultiAnalyst software (Bio-Rad Laboratories).

### 2.6 Quantitative RT-PCR

Differentiated HL-60 cells (3×10^6^) were treated with Cur at different concentrations (1-5 μM) and harvested at various times between 30 minutes to 2 h post-stimulation. Total RNA was extracted from the cells using the Monarch Total RNA Miniprep kit (New England Biolabs, distributed by EuroClone), reverse-transcribed (1 μg, LunaScript RT Supermix 5X; New England Biolabs), and analyzed by quantitative PCR (qPCR) using gene-specific primers (https://www.ncbi.nlm.nih.gov: human NADPH Quinone Oxidoreductase 1, NQO1 Gene ID 1728, Heme Oxygenase HO1 Gene ID: 3162 **Supplementary Table S1**) and Fluocycle IIᵀᴹ SYBR GREEN Master Mix (EuroClone) with an AriaMx instrument (Agilent Technologies, Milan, Italy). The relative levels of gene expression for both target and reference genes were calculated by the 2−^ΔΔCt^ method based on Ct values. The data (n=3 independent experiments performed in quadruplicate) were presented as the mean ± standard deviation (s.d.) of 2−^ΔCt^ values (Livak and Schmittgen, 2001). All mRNA levels were normalized to endogenous GAPDH expression (Gene ID 2597).

### 2.7 Qualitative and quantitative analysis of curcumin in cell extract

For Cur intracellular analysis, High-Performance Liquid Chromatography (HPLC)-grade acetonitrile and methanol were purchased from Merck/Sigma-Aldrich. Glacial acetic acid was obtained from Carlo Erba (Cornaredo, Milan, Italy). The HPLC-grade water (18.2 MW) was prepared using a Millipore Milli-Q purification system (Millipore Corp., Bedford, MA, USA). HL-60 cells (1.5×10^6^) were treated with 1 μM Cur in a complete RPMI cell culture medium to measure Cur levels inside the cells. After 5 minutes, the cells were washed with PBS, then suspended three times in methanol and sonicated. Cur concentrations in the cell extracts were determined using HPLC-Ultraviolet (HPLC–UV) analysis. The analysis was performed with an HP 1110 series HPLC (Agilent, Palo Alto, CA, USA) equipped with a binary pump (G-1312A) and a UV detector (G-1314A). Samples were analyzed using a reverse phase XBridge BEH C18 column (130 Å, 5 mm, 4.6 mm x150 mm) (Waters, Milford, MA, USA) at a flow rate of 1 ml min^-1^. Solvent A was 2% acetic acid, and solvent B was 2% acetic acid in acetonitrile. The gradient for B was as follows: 10% for 3 minutes, from 10% to 20% in 5 minutes, from 20% to 25% in 5 minutes, from 25% to 35% in 5 minutes, isocratic elution (35% B) for 5 minutes, from 35% to 56% in 20 minutes, from 56% to 65% in 7 minutes, from 65% to 95% in 10 minutes. The eluate was monitored at 420 nm.

To confirm the identity of Cur, HPLC eluted peaks were analyzed by Electrospray Ionization multistage Ion Trap Mass Spectrometry (ESI-ITMS^n^) using a Finnigan LCQ DECA XP Max ion trap mass spectrometer (Thermo Finnigan, San Josè, CA, USA), equipped with Xcalibur® system manager data acquisition software (Thermo Finnigan). Mass spectra were recorded from mass-to-charge ratio (m/z) 80 to 1500 in negative ionization mode. The capillary voltage was set at -10 V, the spray voltage was at 3 kV, and the tube lens offset was at -10 V. The capillary temperature was 275°C. Data were acquired in MS, MS/MS, and MS^n^ scanning mode.

The quantification of Cur was carried out using external calibration curves, which were created by repeatedly injecting a fixed volume of Cur standard across a concentration range of 0.01 to 0.1 µg µl^-1^. This was done at three different concentrations, with duplicate injections at each level. All samples were prepared and analyzed in duplicate, and the results were expressed as ng/1.5×10^6^ cells.

### 2.8 Statistical Analysis

A T-Test for students or One-way ANOVA was conducted using GraphPad Prism version 9.5.1 for macOS (GraphPad Software, San Diego, CA, USA), followed by Turkey’s multiple comparisons test. Statistical significance was considered at a p-value of less than 0.05. The results are presented as mean ± standard deviation (s.d.) based on values from independent experiments conducted in duplicate, triplicate, or quadruplicate. Specific p-values were denoted in the figure legends with symbols indicating: * p < 0.05, ** p < 0.01, *** p < 0.001; **** p < 0.0001.

## 3. Results

### 3.1 Characterization of cellular models

Two models of differentiated cells, HL-60 and K-562, were employed to determine the effects of Cur on stress-induced damage caused by Cd or H_2_O_2_. These models were selected to mimic a “normal” immunophenotype. The HL-60 cell line differentiates towards the monocyte line when treated with vitamin D analog EB1089 (VD) (**Figure 1A-D**) (James et al., 1997). In contrast, the K562 cell line differentiates towards the erythroid line when exposed to the natural phytoalexin resveratrol (RESV) (**Figure 1E-H**) (Della Ragione et al., 2003). HL-60 cells were treated at various times (from 24 to 168 h) with EB1089 (**Figure 1A**) at a concentration of 100 nM, while K-562 cells were incubated with 50 μM RESV from 24 to 72 h (**Figure 1E**). Both the growth curves were established using Trypan Blue dye. **Figure 1A** shows that the differentiating agent causes a significant arrest in cell growth starting from 72 h up to 168 h without showing detectable cytotoxicity. The expression of the molecular marker p27^Kip1^ confirmed the effect on cell cycle arrest (**Figure 1B**), as well as the intracellular accumulation of formazan granules in the NBT assay represented the evidence that cells underwent differentiation (**Figure 1D**). In the case of K-562 cells, **Figure 1E** shows that RESV can delay cell proliferation starting from 24 h. This effect was confirmed by Cdc2 downregulation (**Figure 1F**) and emine accumulation in the cytoplasm, as shown with Neutral Red staining (**Figure 1H**) (Russo et al., 2007, Russo and Russo, 2018). **Table 1** summarizes the cytofluorimetric analysis of HL-60 and K-562 cells treated with the differentiating agents, showing that treatment with VD was not toxic and induced an apparent cell cycle arrest in G0/G1 in HL-60 cells (>90%). In K-562 cells treated with RESV, a cell cycle block was observed already after 24 h, with cells accumulating in the G2/M phase (>45%), confirmed also at longer time (data not shown). In particular, RESV-treated K-562 cells are unable to enter into the S phase, and the accumulation of cells in the G2 phase paralleled with a decreased level of Cdc2 protein (**Figure 1F**), as described (Della Ragione et al., 2003). Liu et al. observed a double cell cycle block of K-562 cells when treated with a concentration of RESV higher than 50 μM. These findings suggest that the growth inhibitory effect of RESV on K-562 cells could be due to a cell cycle block at the G1 phase at low doses of the phytoalexin and at the G2/M phase at higher doses (Liu et al., 2010).

**Figure 1.**
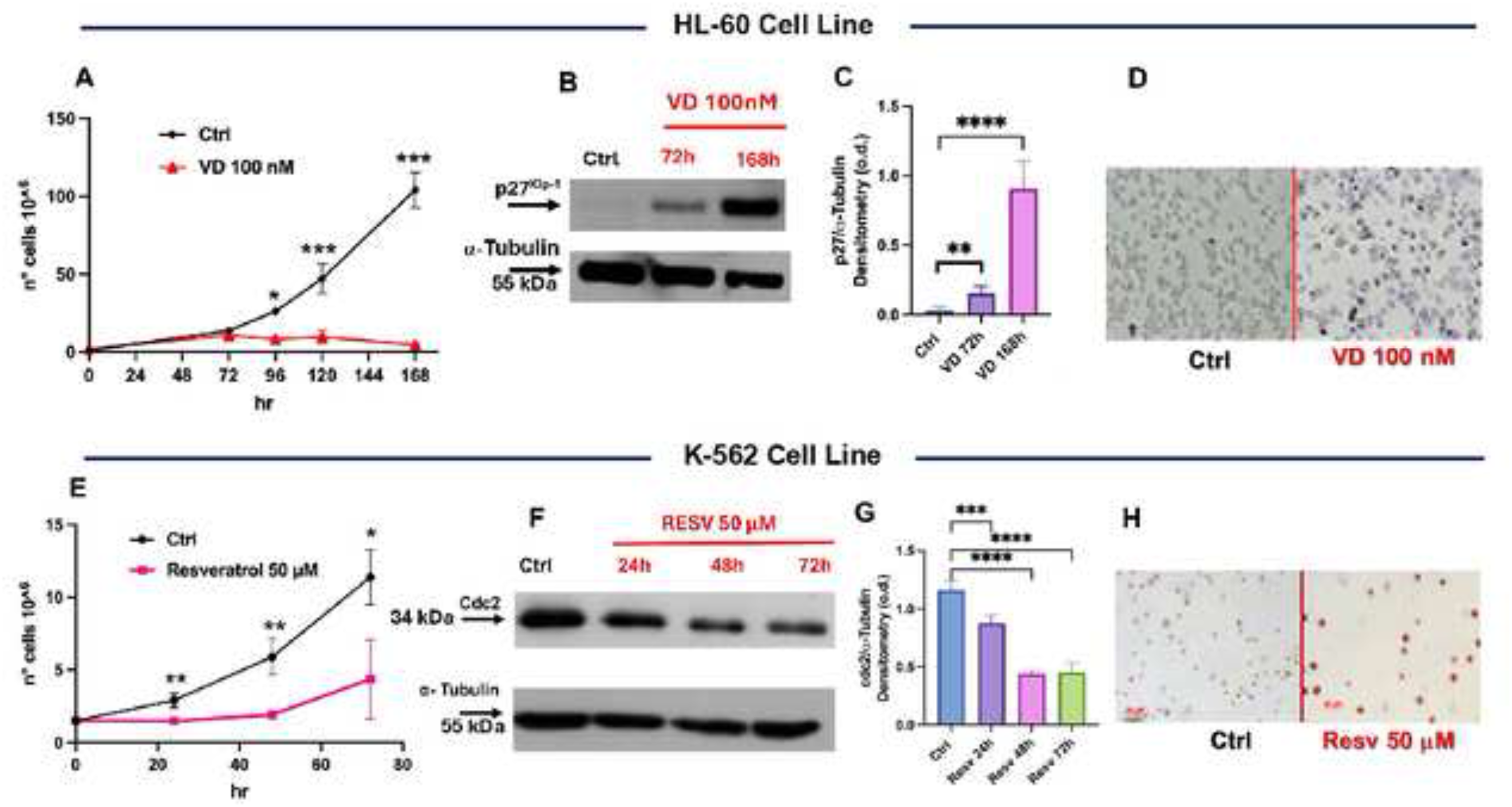
Differentiation of HL-60 and K-562 cells. (**A**) Cell growth curve performed using Trypan blue staining of HL-60 treated with VD and counted at different time points from 72 to 168 h. (**B-C**) Immunoblot evaluation of p27^Kip1^ protein in HL-60 cells post incubation with VD at 72 and 168 h. (**D**) NBT staining of untreated HL-60 cells or treated with VD 96 h. (**E**) Cell growth curve of K-562 cells post incubation with RESV at different time points. (**F-G**) Immunoblot analysis of Cdc2 protein after treatment of K-562 cells with RESV. (**H**) Neutral Red staining of K-562 cells treated with RESV for 72 h. Bars indicate the mean of three duplicate experiments and the standard deviation (± s.d.). The symbols indicate the significance calculated with the T-Test or one-way ANOVA multiple comparison tests using the GraphPad software: *p<0.05 **p<0.01, ***p<0.001 **** p<0.0001.

**Table 1.** Cell cycle analysis in undifferentiated and differentiated HL-60 and K-562 cell lines (percentage of cell cycle distribution)

| Cell Cycle Phase | HL-60<br>(undifferentiated) | HL-60<br>(VD 168 h) | K-562<br>(undifferentiated) | K-562<br>(RESV 24 h) |
| --- | --- | --- | --- | --- |
| **G0/G1** | 36.95±8.5 | 94.2±2.3** | 62±0.8 | 56.7±7.4 |
| **S** | 43.35±9.4 | 8.65±5.4 | 18.2±0.7 | 0 |
| **G2/M** | 9.4±6.3 | 2.5±0.03 | 21.5±0.4 | 46.1±1.4** |

\*\* $p < 0.01$ vs Ctrl cells, T-Test Student, $n = 2$ experiments in duplicate

### 3.2 Pre-incubation with Curcumin protects differentiated cells from Cd-induced toxicity

The Cd concentrations were selected based on the EC_50_ established for both cell lines (**Supplementary Figures S1 and S2**) and optimized to induce a cytotoxicity of about 50% in order to magnify the potential protective effect of Cur. Therefore, CyQuant cell viability assay, reported in **Figure 2A**, shows that in differentiated HL-60 cells, 15 µM Cd reduced cell viability to lower than 50%. However, the pre-treatment for 16 h with low doses of Cur (1 µM) reduced Cd cytotoxicity with a significant protection rate (p<0.05) of approximately 20%. The protective effect of Cur was also evident in the micrograph in **Figure 2B** obtained after cell staining with CyQuant. Here, a significant number of fluorescent cells was present in the combined treatment (bottom right panel) compared to the treated cells with the Cd alone (bottom left panel). **Figure 2C** shows the result of the CyQuant assay in K-562 differentiated cells after pre-incubation with Cur and then treatment with 10 μM Cd for 24 h. In this case, cell toxicity induced by Cd was of about 40%, and Cur pre-treatment was able to significantly protect cells (15%; p<0.05) from death. The CyQuant fluorescence measurement in **Figure 2D** shows representative fields of differentiated K-562 cells treated as indicated to highlight the protective effect of Cur in the combined treatment (bottom left panel).

**Figure 2.**
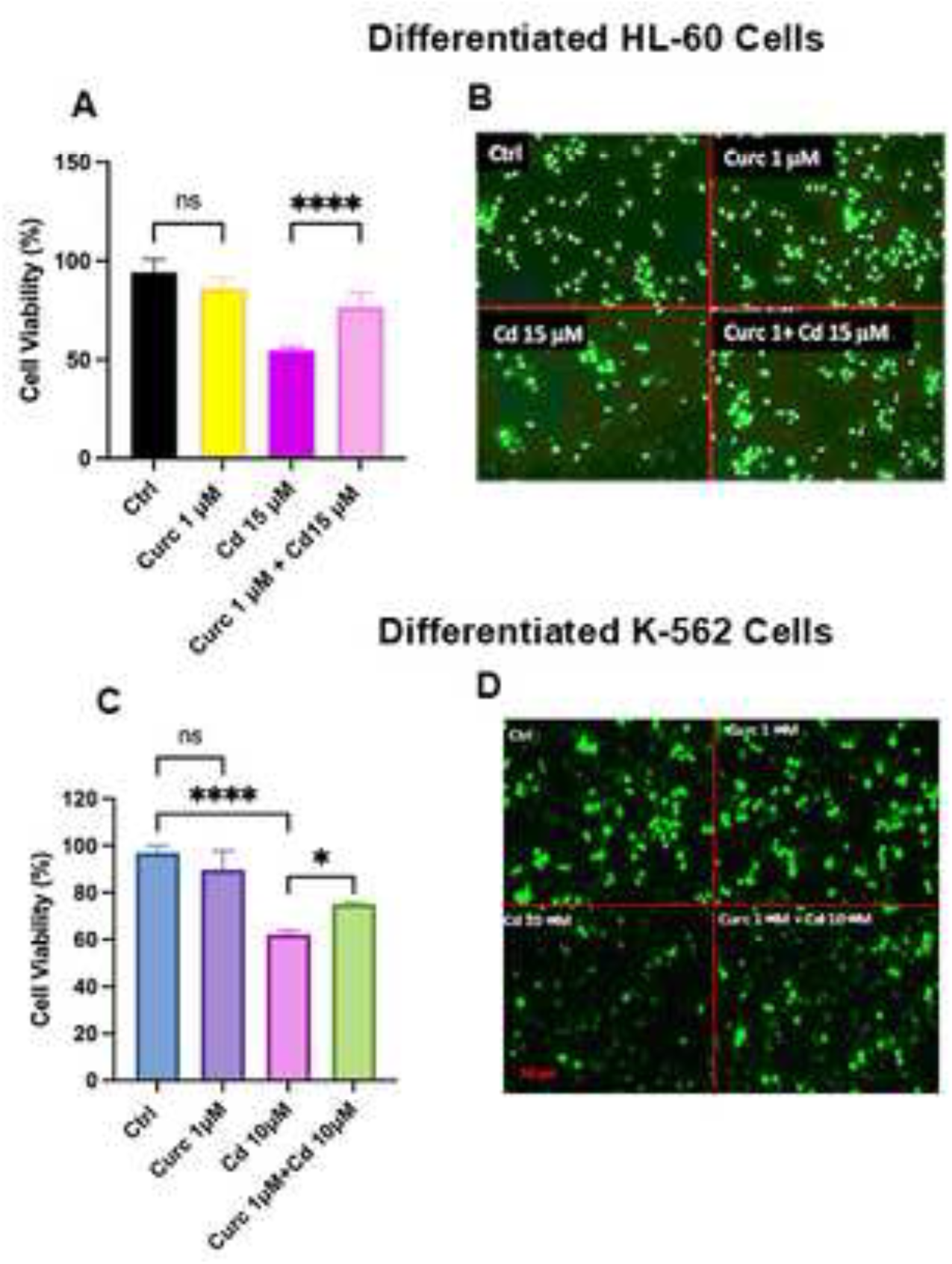
Protective effect of Curcumin against Cd-induced toxicity. HL-60 (panels **A** and B) and K-562 (panels **C** and **D**) cells were incubated with Cur (1 µM) for 16 h and subsequently were treated with Cd for an additional 24 h. At the end of the stimulation, cell viability was measured using the CyQuant assay. The bars in the graphs represent the average of two experiments in quadruplicate (± s.d.). The symbols indicate the significance calculated with one-way ANOVA multiple comparison tests using the GraphPad software: *p<0.05 **p<0.01, ***p<0.001 **** p<0.0001; n.s. not significant. Panels **B** and **D** show representative images of differentiated HL-60 and K562 cells, respectively, following the indicated treatments acquired with the Axiovert 200 fluorescence invertoscope (200X magnification).

### 3.3 Antioxidant Effect of Curcumin on differentiated HL-60 and K-562 cells

To verify whether the cytotoxicity of Cd was related to oxidative damage, the HL-60 and K-562 differentiated cells were incubated with dichlorofluorescein (DCFH) and monochloride bimane (mCB), two fluorochromes capable of measuring the intracellular level of peroxides and GSH, respectively (**Figure 3**). Cur alone, after overnight incubation, was unable to modify intracellular peroxide significantly (**Figure 3AB**). However, at the same time point, the molecule increased GSH intracellular levels in both cell lines (**Figure 3CD**). Moreover, as expected, Cd induced significant oxidative stress in differentiated K-562 and HL-60 cells, resulting the increased levels of ROS of about 30% and 15%, respectively (**Figure 3AB**), which paralleled with lower intracellular GSH (**Figure 3CD**), one of the leading systems capable of controlling the intracellular redox state.

**Figure 3.**
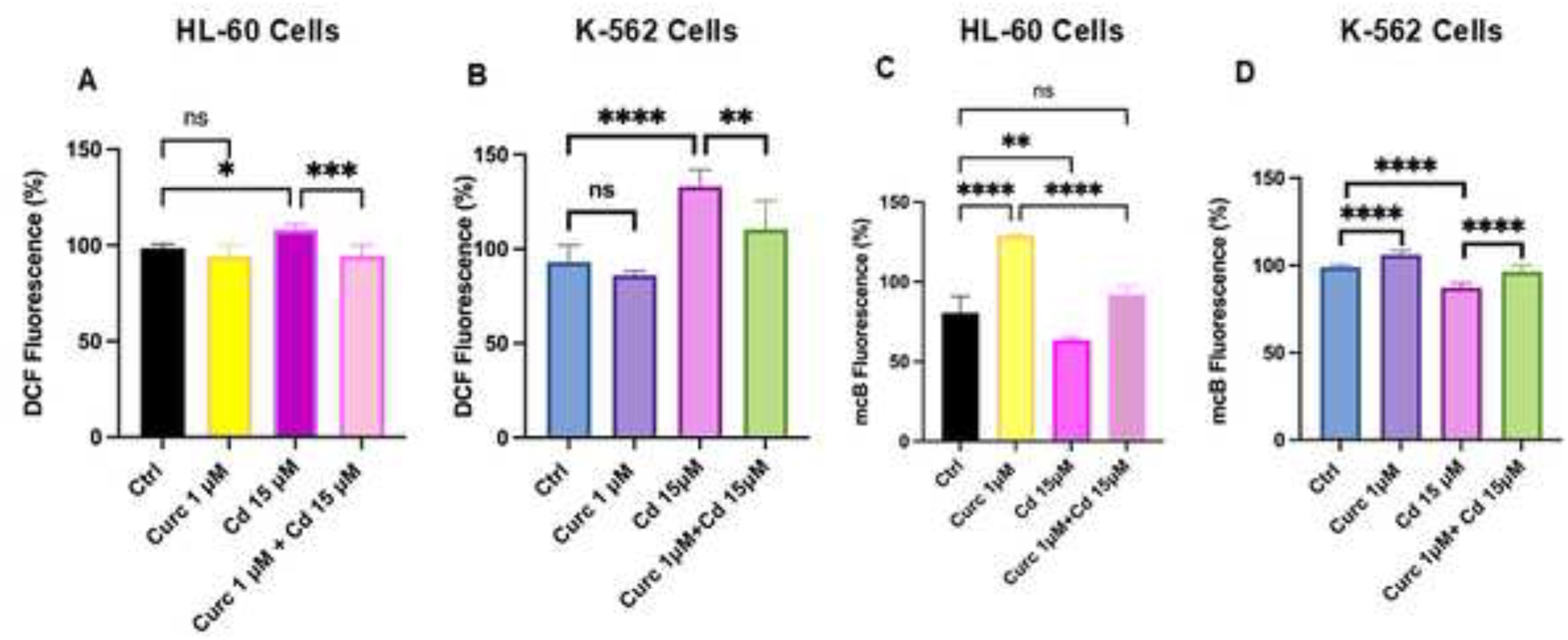
Antioxidant effect of Curcumin in differentiated cells. Differentiated cells were preincubated with 1 μM Cur for 16 h and treated with Cd at the indicated concentration for 30 minutes in HL-60 cells (panel **A**) and 4 h in K-562 cells (panel **B**). Intracellular peroxides were measured with a CM-DCFDA probe and indicated as a percentage of DCF fluorescence. GSH levels (panels **C** and **D**) were fluorimetrically measured using an mCB probe after pre-incubation of cells with Cur and then treated with the indicated concentration of Cd for 4 h. The bars in the graphs represent the average of two experiments in quadruplicate (± s.d.). The symbols indicate the significance calculated with one-way ANOVA multiple comparison tests: *=p<0.05 **=p<0.01, ***=p<0.001 **** =p<0.0001; n.s. not significant.

Finally, when cells were pre-incubated with Cur and treated with Cd, intracellular peroxide levels were significantly reduced with respect to Cd treatment in both cell lines (**Figure 3AB**). The antioxidant effect observed after Cur pre-incubation was confirmed by GSH levels that returned to the basal levels in the combined treatment (**Figure 3CD**).

### 3.4 Curcumin protects differentiated HL-60 and K-562 cells against H_2_O_2_ induced toxicity

To reinforce the hypothesis that Cur treatment could modify redox status in both cell lines, rendering them more responsive to subsequent oxidative stress, we treated differentiated HL-60 and K-562 cells with different doses of H_2_O_2_. CyQuant viability assay was employed to determine the ideal concentration of H_2_O_2_ for the subsequent cell protection experiments (**Supplementary Figures S2**). In **Figure 4A**, 20 μM H_2_O_2_ was applied to evaluate the protective effect of Cur pre-incubation on differentiated HL-60 cell viability. The protection was of about 30% with respect to the single treatment with H_2_O_2_. Pre-incubation with Cur in differentiated K-562 cells showed the same trend even with a higher concentration of H_2_O_2_ (50 μM), which reduced cell viability by about 40% (**Figure 4B**). The presence of Cur almost completely rescued the H_2_O_2_-induced toxicity in K-562 (**Figure 4B**). Measuring intracellular peroxide after overnight pre-incubation with Cur and then damaging cells for 30 minutes with H_2_O_2_, confirmed its protective capacity against oxidative stress in both differentiated cell lines (**Figure 4CD**). In particular, Cur preincubation for only 2 h in differentiated K-562 cells was able to exert significant protection against ROS increase (<20% production; **Figure 4D**).

**Figure 4.**
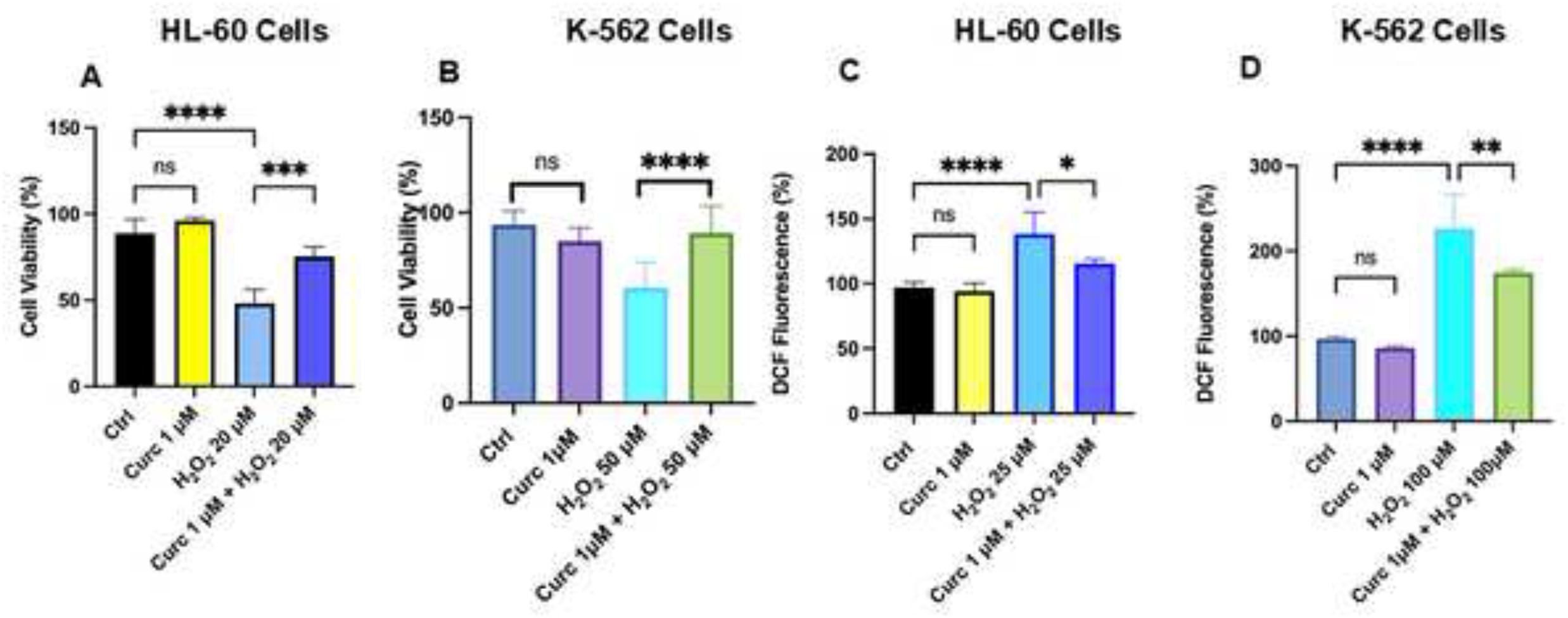
Curcumin protects differentiated cells from H_2_O_2_ damage. Cells were preincubated with 1 μM Cur for 16 h and then treated with H_2_O_2_ at the indicated concentration for 24 h in differentiated HL-60 (panel **A**) and in K-562 (panel **B**) cells. Cell viability was evaluated with CyQuant assay. In panel **C**, the amount of intracellular peroxides was measured using a CM-DCFDA probe after overnight incubation with Cur followed by 30 minutes treatment with H_2_O_2_ in differentiated HL-60 cells. Similarly, after 16 h of Cur incubation in differentiated K-562 cells, the amount of peroxides was measured after 30 minutes and reported as a percentage of DCF fluorescence (panel **D**). The bars in the graphs represent the average of two experiments in quadruplicate, with the standard deviation represented by ± s.d. The symbols used indicate significance calculated with one-way ANOVA multiple comparison tests: *=p<0.05, **=p<0.01, ***=p<0.001, ****=p<0.0001; n.s. indicates not significant.

### 3.5 Curcumin induces a mild pro-oxidant effect in differentiatedHL-60 and K-562 cells

We investigated the redox status of differentiated HL-60 and K-562 cells soon after Cur incubation using a specific probe for superoxide anion (DHE) after 40 minutes of Cur treatment in both cells. **Figure 5A** shows that Cur significantly increased O^-^ intracellular levels in differentiated HL-60 cells (> 2 fold). Similarly, the pro-oxidant effect of Cur was also confirmed in differentiated K-562 cells (>40%) (**Figure 5B**).

**Figure 5.**
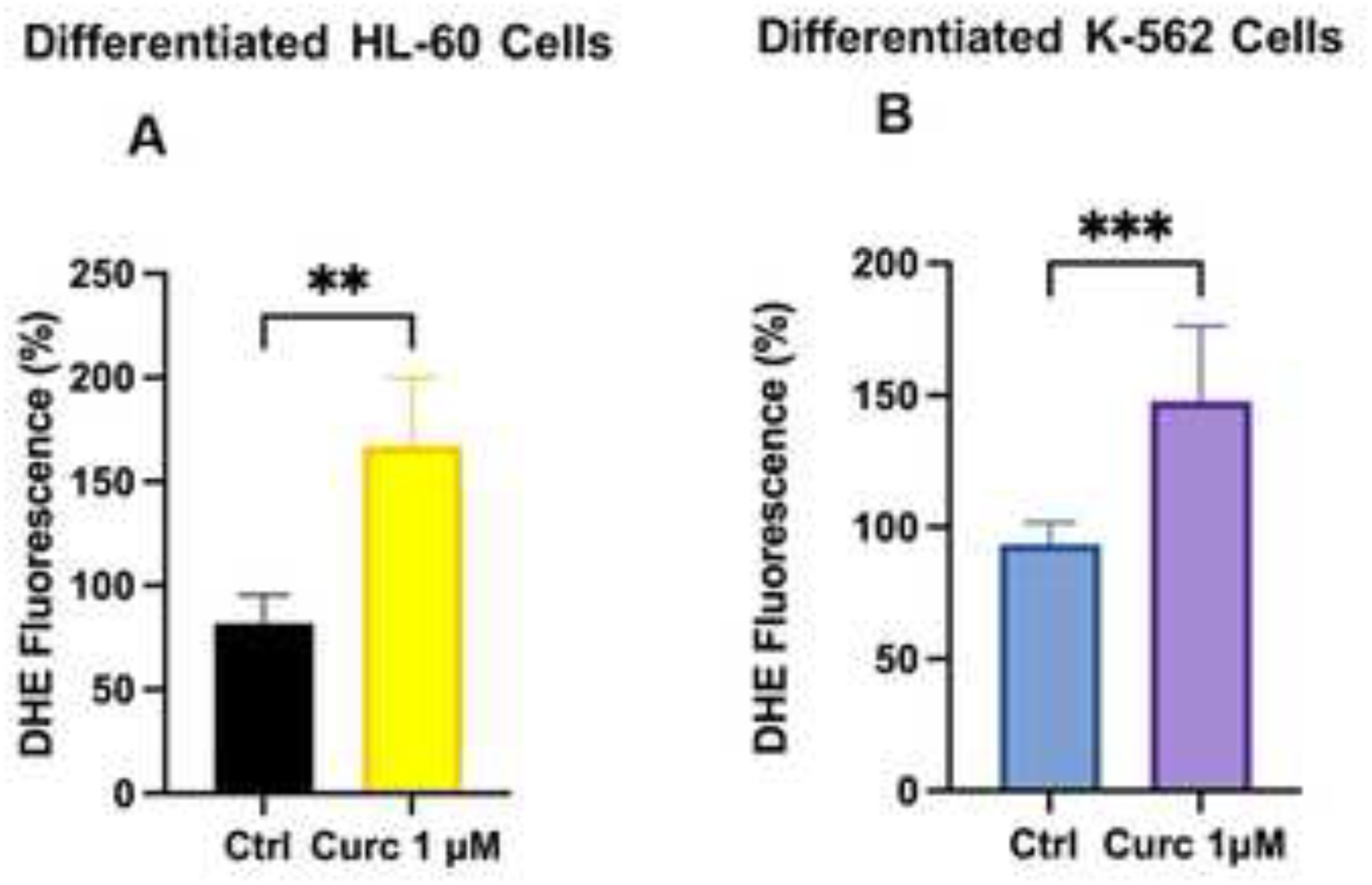
Curcumin increases intracellular superoxide anion. Differentiated HL-60 (panle **A**) and K-562 (panel B) cells were incubated for 40 minutes with 1 μM Cur. A DHE probe was used to fluorimetrically measure intracellular levels of superoxide anion expressed as a percentage of DHE fluorescence. The bars in the graphs represent the average of two experiments in quadruplicate, with the standard deviation represented by ± s.d. The symbols used indicate significance calculated with Student’s T-Test (panel **A**) or one-way ANOVA multiple comparison tests (panel **B**): *=p<0.05, **=p<0.01, ***=p<0.001, ****=p<0.0001; n.s. indicates not significant.

### 3.6 Curcumin uptake in HL-60 cell line

Given the short half-life of Cur in water, it is challenging to discern whether the biological effects observed *in vitro* were caused by Cur itself or its breakdown products, which makes difficult to fully understand the mechanisms of Cur biological actions (Nimiya et al., 2016). Therefore, we developed a method to extract curcumin from HL-60 cells shortly after adding it to the cell culture medium (5 minutes) (Russo et al., 2017). In Table 2, it is shown that free Cur was detected in cells after 5 minutes of incubation. However, it was quickly degraded over time (from 1 to 15 h of incubation; data not shown). This finding suggests that even at low levels, Cur can be rapidly absorbed in HL-60 cells, where it can potentially exert antioxidant or pro-oxidant effects against the cytotoxicity of Cd and H_2_O_2_.

**Table 2.** Intracellular levels of Curcumin in HL-60 cell.

| Ctrl (DMSO 0.1%) | Treatment | Curcumin [$1 \mu\text{M}$]* |
| --- | --- | --- |
| n.d. | 5 minutes | 11.81±4.52 |

\*Numbers indicate Curcumin as $\text{ng}/1.5 \times 10^6$ cells ± s.d.

### 3.7 Curcumin antioxidant activity involves Nrf2 transcription factor

Understanding the mechanism by which Cur shields cells against oxidative damage caused by Cd or hydrogen peroxide is crucial. To achieve this goal, we investigated the role of Nrf2, a pivotal transcription factor in the antioxidant response of nucleated cells. Polyphenols, such as Cur, have been linked to regulating the redox status of cells by targeting Nrf2 (Chun et al., 2021). Therefore, we assessed the presence of Nrf2 protein in the nuclear compartment and measured by qRT-PCR the levels of two canonical Nrf2-regulated transcripts, HO-1 (heme oxygenase-1) and NQO1 (NAD(P)H quinone dehydrogenase 1), in differentiated HL-60 cells.

The immunoblot experiment in **Figure 6A** shows that after 2 h of Cur treatment, the expression of Nrf2 protein in the nuclear compartment increased. Nrf2-dependent transcript levels were measured by qRT-PCR on RNA extracted from differentiated HL-60 cells after 30, 60, and 120 minutes of incubation with 1 μM Cur. The time course for qRT-PCR analysis was carefully chosen based on the consolidated data regarding the nuclear translocation of Nrf2 (2 h) (**Figure 6A**), ROS protective effects of Cur after 16 h of treatment (**Figure 3A** and **4A**), and changes in the intracellular ROS levels after 40 minutes (**Figure 5A**). The results show a significant increase in the HO gene transcript after 2 h of incubation at low doses of Cur (1 μM) (>50%; **Figure 6C**). Using a higher dose of Cur (5 μM), the levels of mRNA of the HO gene almost doubled (**Supplementary Figure S3**). Similarly, NQO1 transcript significantly increased (>50%) after 2 h of treatment with 1 μM Cur (**Figure 6D**). These data indicate that low doses of Cur in differentiated HL-60 cells can activate the transcription of genes under the control of the transcriptional factor Nrf2 with the result of counteracting oxidative damage.

**Figure 6.**
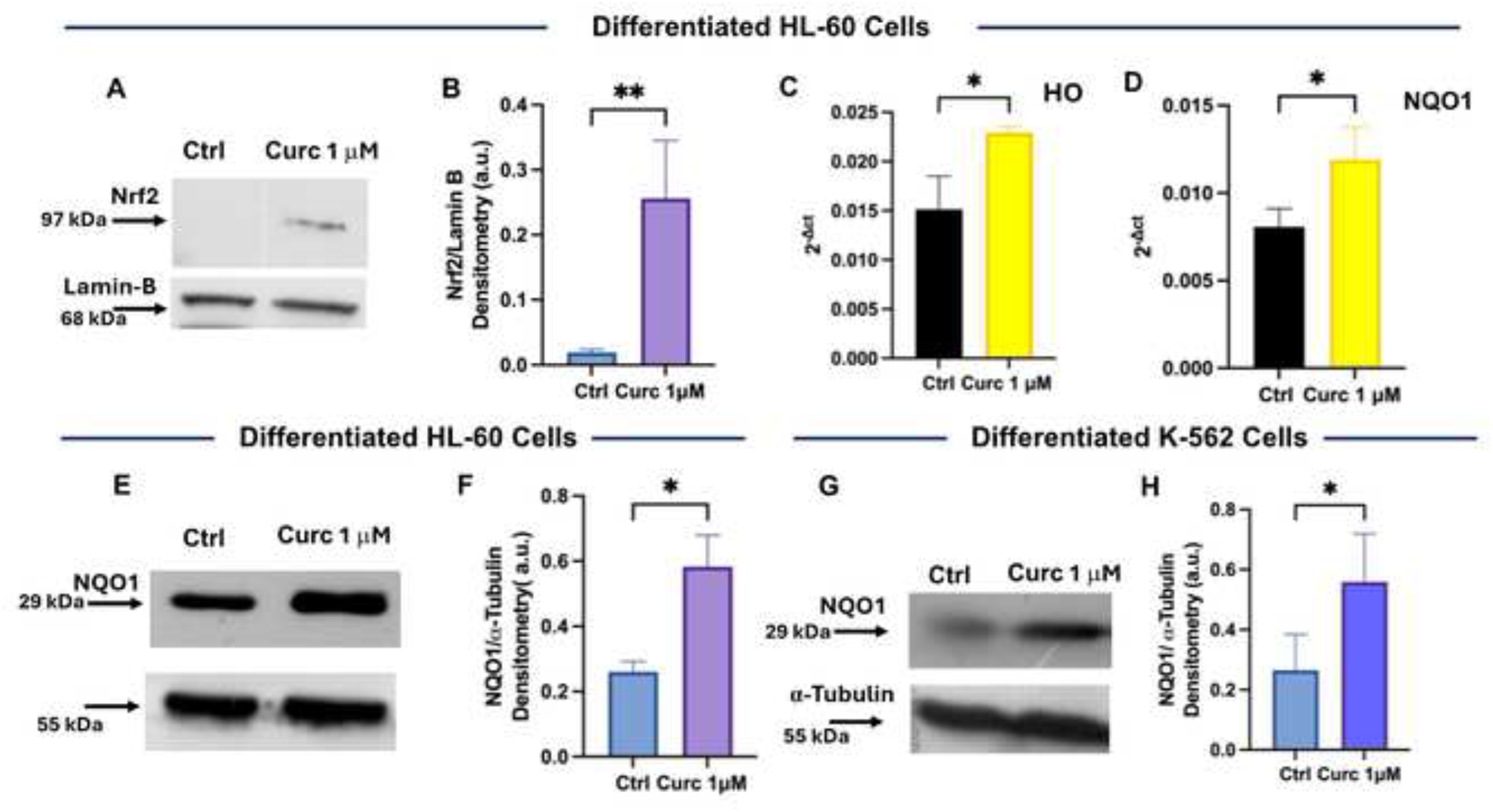
Curcumin activates Nrf2-dependent antioxidant response. Differentiated HL-60 cells (panel **A**) were incubated with 0.1% DMSO or 1 μM Cur for 2 h, and the nuclear fraction was loaded on SDS-PAGE. Immunoblot analysis was performed to mesure Nrf2 expression. Band quantification was achieved with Multi-Analyst software by measuring the ratio of Nrf2/Lamin-B densitometry (panel **B**). In panels **C** and **D**, differentiated HL-60 cells were incubated with 0.1% DMSO (Ctrl) or 1 µM Cur for 2 h. The RNA was extracted and reverse transcribed, and the specific sequences for the transcripts were amplified and quantified by qRT-PCR. The bars indicate the average of three quadruplicate experiments with values expressed as 2^-ΔCt^ ± s.d. Symbols indicate significance calculated with ANOVA: * = p<0.05; ** = p<0.01. Differentiated HL-60 (panels **E** and **F**) or K-562 (panels **G** and **H**) cells were incubated with 0.1% DMSO or 1 μM Cur for 16 h or 2 h, respectively. Immunoblots were performed using anti-NQO1 antibody. Bands quantification was achieved with Multi-Analyst software (BioRad) by measuring the ratio of NQO1/α-Tubulin densitometry. The bars indicate the average of 2 experiments ± s.d, and symbols indicate significance calculated with Student’s T-Test : * = p<0.05.

To confirm the accuracy of the data obtained from the transcripts, we exposed differentiated HL-60 and K-562 cells to 1 μM Cur for 16 h and 2 h, respectively, and measured the expression of NQO1 protein. Immunoblot analyses are presented in **Figure 6EF** for HL-60 cells and Figure **6FG** for K-562 cells, respectively. The results suggest that Cur, by increasing the expression of NQO1 by over 2-fold, may protect cells from the subsequent oxidative insult generated by treatments with oxidative agents (e.g., Cd or H_2_O_2_).

### 3.8 Curcumin induces protective autophagy in differentiated K-562 and HL-60 cells

Autophagy is a mechanism of cellular response to damage caused by ROS that has been extensively studied (Mizushima and Levine, 2020). We looked at markers of autophagy in differentiated K-562 and HL-60 cells after treatment with Cur for 2 h. Therefore, we measured changes in the expression of p62^SQSTM1^, a multifunctional protein involved in signal transduction pathways linking autophagy and the Keap1–Nrf2 pathway (Zhong et al., 2016). In particular, the expression of p62^SQSTM1^ increased with the accumulation of autophagosomes (**Figure 7A-D**) (Moosavi et al., 2018). Its expression increased of 2.5-fold after 2 h of treatment with 1 μM Cur in differentiated K-562 cells (**Figure 7AB**). A less pronounced but still significant increased expression of p62^SQSTM1^ was also detectable in differentiated HL-60 cells (>40%; **Figure 7CD**).

**Figure 7.**
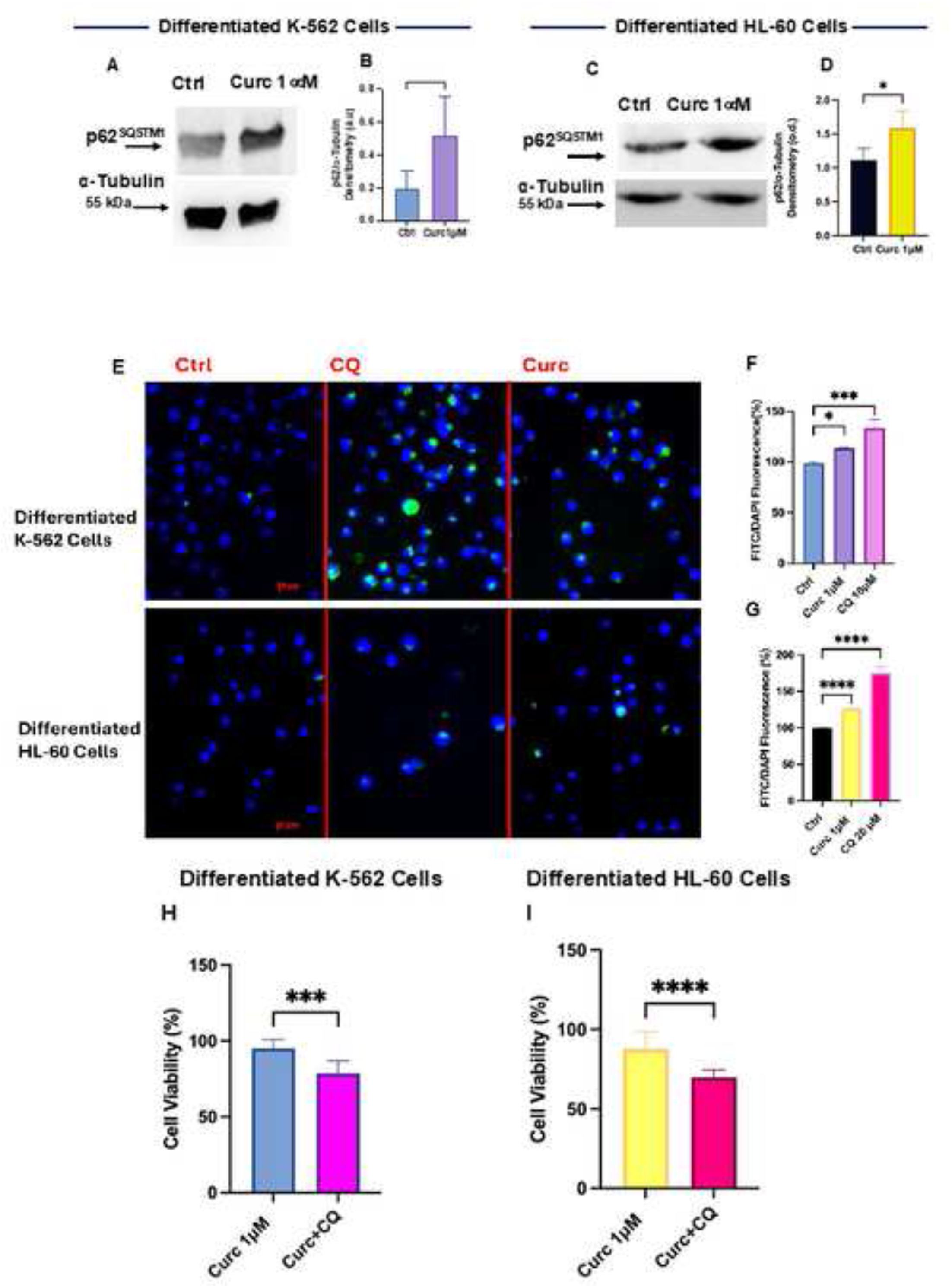
Curcumin modifies autophagic flux in differentiated K-562 and HL-60 cells. Differentiated K-562 (panels **A** and **B**) and HL-60 (panels **C** and **D**) cells were incubated for 2 h with 1 μM Cur. Subsequently, cellular homogenates were employed for Immunoblot analysis to assess the expression of p62^SQSTM1^. Quantification of protein bands is shown in the histograms (panels **B** and **D**) and was achieved by Multi-Analyst software by measuring the ratio of p62^SQSTM1^/α-Tubulin. In panels **E**, **F** and **G**, Cyto ID assay was applied to fluorimetrically quantify intracellular autophagosomes at the end of Cur or CQ treatment as described in the Materials and Methods section. Panel **E** shows micrographs of representative fields of K-562 and HL-60 cells using an inverted fluorescence microscope (Axiovert 200 Zeiss, 400x magnification) after Cyto-ID quantification. In panels **H** and **I**, Cy-Quant viability assay was performed to examine the role of Cur-induced autophagy on differentiated K-562 (panel **H**) or HL-60 (panel **I**) cells treated for 24 h with Cur alone or after pre-incubation with CQ. The bars indicate the average of two experiments in quadruplicate ± s.d, and symbols indicate significance calculated with Student’s T-Test: * = p<0.05; *** = p<0.001; **** = p<0.0001 with ****.

To confirm that Cur can alter autophagic flux in differentiated K-562 and HL-60 cells, we quantified the intracellular autophagosomes using Chloroquine (CQ) as a positive control, an inducer of autophagosome accumulation into the cytoplasm. The histogram in **Figure 7F** shows the specific quantification of intracellular autophagosomes in K-562 cells with fluorescent probe Cyto-ID (green dots), which was normalized with respect to Hoechst nuclear dye (blue stain in micrograph in **Figure 7E**). In differentiated K-562 cells, Cur increased by 15% the number of cytoplasmic autophagosomes with respect to DMSO-treated cells, while for CQ, the increase was >30% (**Figure 7F**). The same approach and quantification were applied to differentiated HL-60 cells (compare **Figure 7G** with lower panels in **Figure 7E**), where Cur increased of about 40% the number of autophagosomes, while CQ, as expected, was more effective with an enhancement of autophagosomal number > 80%.

Autophagy is a complex process with multiple modes of action, including cytoprotective versus cytotoxic functions (Wei et al., 2024). To prove that Cur induced a protective form of autophagy in our cellular models, we conducted a cell viability experiment. We preincubated cells with CQ, which blocked the autophagic flux, and then added Cur to evaluate cell viability with CyQuant dye (**Figure 7HI**). After 24 h of either Cur treatment or the combined treatment, we found that Cur induced a protective form of autophagy because cell viability decreased significantly with respect to Cur single-treatment (**Figure 7HI**). To better comprehend this result, it should be considered that when a form of protective autophagy is activated, the autophagic flux is modified to recycle and repair damaged cellular structures. This process is no longer active if the autophagic flux is blocked and cytotoxicity occurs.

## 4. Discussion

Several studies in the literature have evaluated the protective role of Cur-induced autophagy against oxidative stress in different cellular models. Among these, a recent study demonstrates that Cur can protect against pro-oxidant and inflammatory damage induced by arsenic in mouse models by activating an autophagic process dependent on the transcription factor TFEB (Xu et al., 2024). In another study, it was demonstrated that Cur attenuates the toxicity of diphenyl diphosphate, a toxic derivative of the processing of plastic materials, through the modulation of autophagic flux and the transcriptional factor Nrf2, responsible for the protein-dependent antioxidant and anti-apoptotic response p53 (Wei et al., 2024).

The present work strongly supports the existing literature and demonstrates that even at low doses, Cur can stimulate a protective homeostatic response that involves autophagy at the cellular and probably tissue level in response to external stressors and environmental pollutants (Calabrese et al., 2007, Lin et al., 2019).

As mentioned in the Introduction, autophagy can have different roles during carcinogenesis, sometimes opposite, protective, or lethal depending on the stage of tumor progression, reducing the effects of cellular stresses related to the genesis of tumors or, in advanced stages, promoting the unwanted survival of malignant cells (Russo and Russo, 2018). As regards the lethal or cytotoxic effect, autophagy can enhance various forms of cell death even if the molecular mechanisms through which it exerts a tumor-suppressive role are still poorly understood (Russo et al., 2022). In the case of Cur, the activation of protective autophagy in differentiated cells is undoubtedly a desirable process as it protects them from the damage caused by ROS on the DNA, preventing genomic instability and the consequent activation of oncogenes related to tumor progression. Furthermore, the activation of autophagy under stress conditions also avoids the establishment of inflammatory processes and the resultant damage to the surrounding tissue and, over time, the progression toward the metastatic phenotype.

As regards the mechanism of action, like other polyphenols, such as EGCG (epigallocatechin gallate) (Lee et al., 2024), the activation of protective autophagy by Cur may be related to the antioxidant response dependent on Nrf2. In fact, in recent papers, it has been demonstrated that EGCG behaves differently in normal cells compared to neoplastic ones by positively regulating the p62/Nrf2/Keap1-dependent signaling pathway in normal cells and inhibiting it in tumor cells, selectively inducing cell death by apoptosis (Lee et al., 2024).

In the present study, the levels of the multifunctional protein p62^SQSTM1^ were found to be higher in Cur-treated cells. Cur-induced p62^SQSTM1^ accumulation in myeloid cells could promote Nrf2 translocation to the nucleus, as suggested by the increased activation of NQO1 and HO-1 at the same experimental time points and this mechanism is potentially in agreement with an interaction between Cur or its metabolites with cytoplasmic Keap1 with negatively regulates Nrf2 activation (Komatsu et al., 2010).

To determine the impact of perturbation of cellular homeostasis on cell survival or death, several factors come into play, including the variation of the intracellular redox state. In the cellular models used in this study, a low dose of Cur was found to be rapidly absorbed by cells within 5 minutes (**Table 2**), and the molecule itself, or its metabolites or oxidized products, were able to increase the expression of HO and NQO1 transcripts and proteins after 2 h or overnight treatment, respectively. Furthermore, after 4 h, Cur also increased the levels of intracellular GSH, thus enhancing cellular defenses against exposure to stress factors. In this study, stress factors were experimentally induced by treating cells with Cd or H_2_O_2_, both of which were found to lower intracellular GSH levels in the cellular models.

In biology and medicine, hormesis denotes the adaptive response of cells and organisms to moderate, typically intermittent, stress (Calabrese et al., 2007, Calabrese et al., 2023). The low concentration of Cur employed in the present study is compatible with a hormetic dose that “directly” or “indirectly” can trigger an adaptive cellular response by acting on the Nrf2/Keap1 system (Calabrese et al., 2007, Russo et al., 2024). Our hypothesis is that the presence of Cur, even at low concentrations, induces a moderate cellular pro-oxidant response ((Nimiya et al., 2016) and **Figure 5**), not sufficient to induce cellular damage but enough to activate protective autophagy and the antioxidant response dependent on p62^SQSTM1^/Nrf2/Keap1, as suggested by the increased expression of GSH HO and NQO1.

Several studies have suggested that certain phytochemicals, including sulforaphane, quercetin, and Cur, can act as electrophiles that can bind to specific cysteines (Cys^151^) on Keap1 (Shin et al., 2020). This binding can result in a conformational change of the E3 ubiquitin ligase, Keap-1, leading to a decrease in its inhibitory activity and an increase in the stability of Nrf2. This view is in agreement with the results presented here (**Figure 5**), showing the pro-oxidant behavior of Cur. An intrinsic limitation of our study, as mentioned in the Introduction section, is represented by the low bioavailability of Cur, which is a very unstable molecule with rapid clearance (**Table 2**) and possesses toxical potential, resulting in its categorization as a pan-assay interference compound (PAIN) (Nelson et al., 2017), although the low concentrations employed in the present work strongly limits this eventuality. A thorough review of the literature found that there are very few double-blind human clinical trials that demonstrate the therapeutic effect of Cur (Trial NCT00365209) (Russo et al., 2024). Thus, it is crucial to ascertain that Cur possesses real therapeutic potential rather than being a mere “PAIN”. To achieve this goal, appropriate *in vivo* studies with animal models and well-designed, randomized human clinical trials are required. In this study, we chose a low concentration of Cur *to imitate* the low plasmatic concentration of phytochemicals after ingesting a nutraceutical or a “pill” enriched with Cur. We are confident that our findings are indicative of the biological activity of Cur. In fact, Cur is present inside cells soon after (5 minutes) of its incubation (**Table 2**), produces mild oxidative stress after 40 minutes (**Figure 5**), and induces the nuclear translocation of Nrf2 and the transcription of antioxidant response genes after 2 h (NQO1, HO, **Figure 6)**.

Although the low concentration of Cur is an intrinsic limitation due to the chemical instability of the molecule in an aqueous solution (Cheng et al., 2001, Jacob et al., 2022), this concentration is very close to those used in our study and indicates potential biological/protective effects in the organism that will be evaluated carefully in future *in vivo* studies.

The use of nanoparticle-based drug delivery systems presents a promising solution for the challenges in enhancing the effectiveness of Cur in various therapeutic/nutraceutical applications (Jacob et al., 2022).

In summary, low-dose Cur has been shown to be effective in boosting the cell’s defense mechanism against stress factors. Its moderate pro-oxidant effect activates protective autophagy and the antioxidant response through the p62^SQSTM1^ protein and/or the Keap-1/Nrf2 signaling pathway. These findings can help develop new strategies to counteract the damaging effects of ROS and help improve the survival of cells in stressful environments (**Figure 8**).

**Figure 8.**
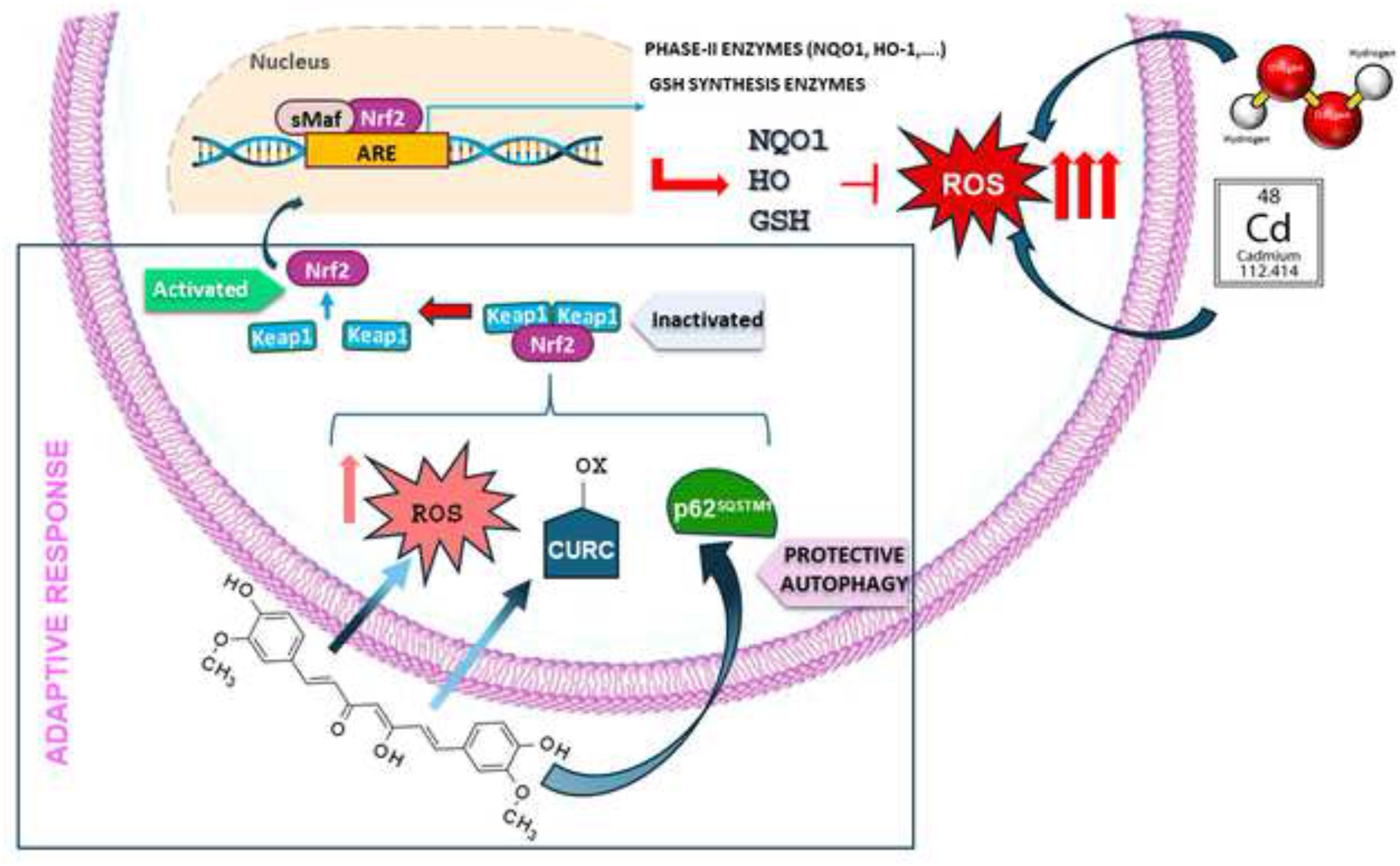
Preventive effects of Curcumin against oxidative stress induced by Cadmium. The proposed mechanism of action of low doses of Cur in safeguarding differentiated K-562 and HL-60 cells from oxidative stress induced by Cd and H2O2 involves the internalization of Cur within cells. The metabolites or oxidized form of Cur activate an adaptive response that induces protective autophagy through the p62SQSTM1 protein. Additionally, Cur promotes an antioxidant response that depends on the Keap-1/Nrf2 signaling pathway by enhancing the levelsof GSH, HO, and NQO1.

The results of this paper can be interpreted as a *proof-of-concept* of the ability of curcumin to preserve cellular homeostasis at nutritional dosages, offering protection from damage caused by exposure to toxic agents present in the environment. This finding has significant implications for the field of phytochemicals, as it suggests that these compounds may have unique mechanisms of action that could be exploited for therapeutic purposes. Further investigation is essential to fully grasp the implications of these findings, yet the potential benefits are unmistakable. In fact, these results lay the foundations for future studies aimed both at exploring the extension of this mechanism to other bioactive compounds present in the diet and at promoting the design of studies on animal models and in humans for the prevention of chronic and degenerative diseases.

## 5. Conclusions

This study demonstrates for the first time that low doses of Cur can protect cells against Cd toxicity. This protection is attributed to the fast uptake of Cur into cells, which results in mild intracellular oxidative stress and an “indirect” antioxidant response. This response relies on Nrf2/ARE transcripts such as HO and NQO1(He et al., 2023, Liu et al., 2017). Additionally, Cur increases intracellular GSH and activates a protective form of autophagy (Tang et al., 2022), an essential process linked to maintaining cellular homeostasis against internal and external stressors (Musial et al., 2021, Russo and Russo, 2018). These findings could help develop Cur-enriched supplements to be tested in interventional studies aimed at protecting people living in areas contaminated by heavy metals.

## Acknowledgements

We thank all the Partners and colleagues of the EcoNutraPrevention project, which inspired the scientific rationale of the present study.

## Author statement

Conceptualization, **Maria Russo** and **Gian Luigi Russo**.; methodology, **Maria Russo, Annamaria Di Giacomo, Federica Fiore, Carmela Spagnuolo, Paola Minasi, Virginia Carbone**; validation, **Maria Russo, Annamaria Di Giacomo, Federica Fiore, Carmela Spagnuolo, Paola Minasi, Virginia Carbone**; Investigation, **Maria Russo, Annamaria Di Giacomo, Federica Fiore, Carmela Spagnuolo, Paola Minasi, Virginia Carbone**; data curation, **Maria Russo**; writing— original draft preparation, **Maria Russo**; writing—review and editing, **Maria Russo**, **Gian Luigi Russo**.; visualization, **Maria Russo, Annamaria Di Giacomo, Federica Fiore, Carmela Spagnuolo,** supervision,. **Maria Russo** and **Gian Luigi Russo**; funding acquisition, **Gian Luigi Russo.**

All authors have read and agreed to the published version of the manuscript.”

## Supplementary Figures and Table

**Figure S1.**
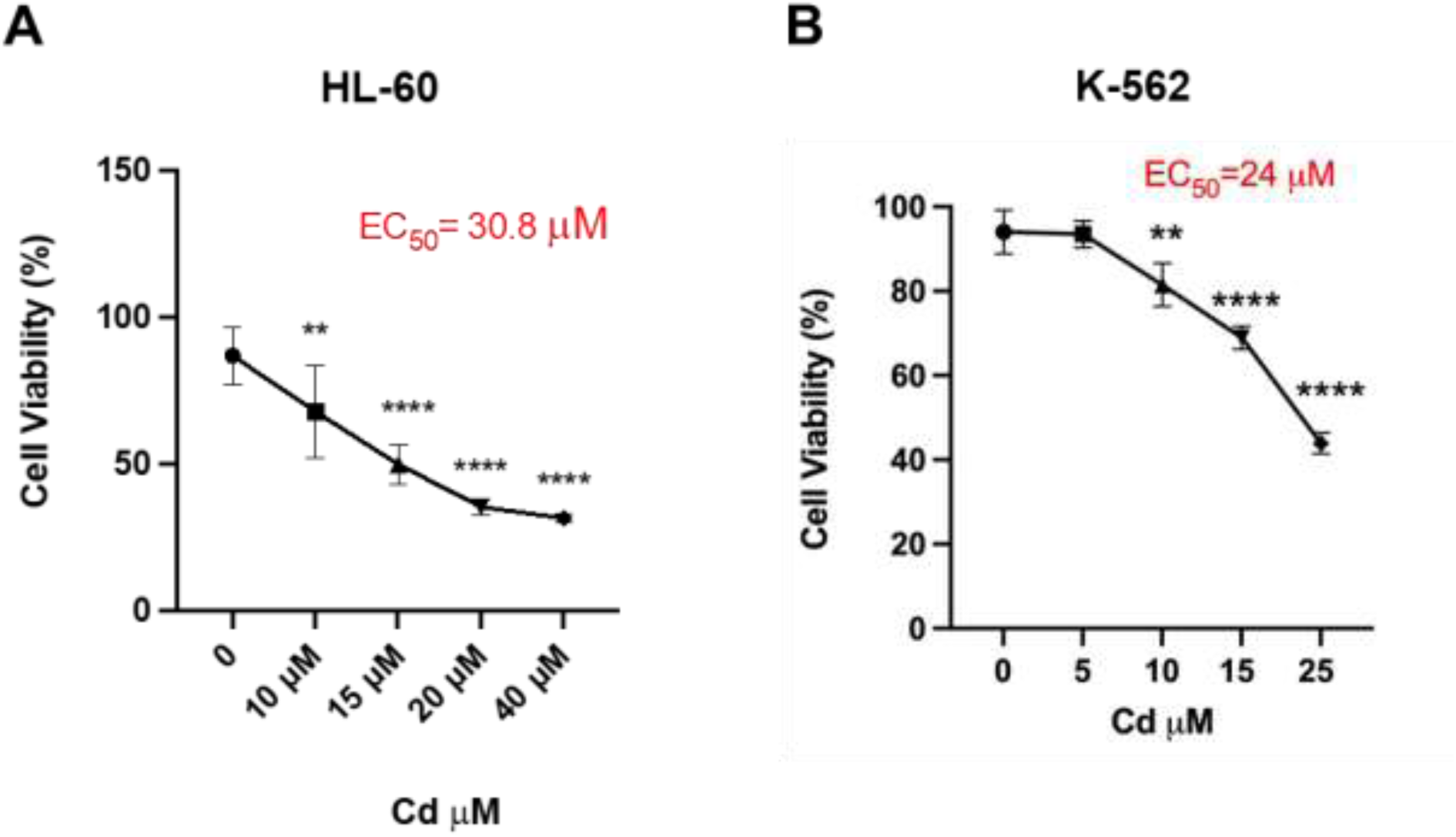
Dose-effect curve of Cadmium cytotoxicity in HL60 and K-562 differentiated cell lines. Differentiated HL-60 (panel A) and K-562 (panel B) cells were incubated with different concentrations of Cd for 24 h. At the end of the stimulation, cell viability was measured using the CyQuant assay. The bars in the graphs represent the average of two experiments in quadruplicate (± s.d.). The symbols indicate the significance calculated with one-way ANOVA multiple comparison tests using the GraphPad software: **p<0.01, ***p<0.001 **** p<0.0001.

**Figure S2.**
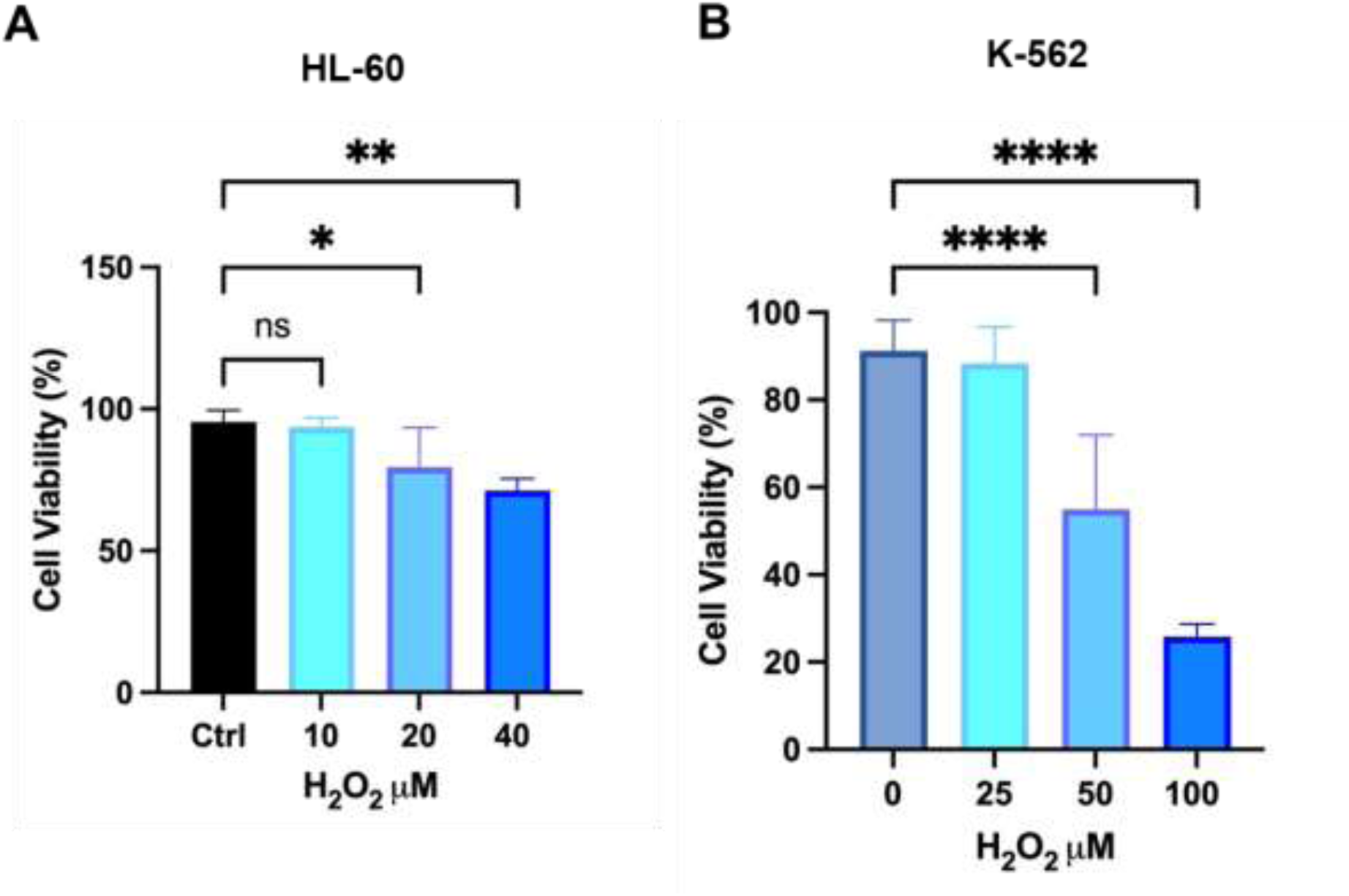
Dose-effect curve of H_2_O_2_ in HL-60 and K-562 differentiated cells. Differentiated HL-60 (panel A) and K-562 (panel B) cells were incubated were incubated with different concentrations of H_2_O_2_ for 24 h. At the end of the stimulation, cell viability was measured using the CyQuant assay. The bars in the graphs represent the average of two experiments in quadruplicate (± s.d.). The symbols indicate the significance calculated with one-way ANOVA multiple comparison tests using the GraphPad software: *p<0.05 **p<0.01, **** p<0.0001.

**Figure S3.**
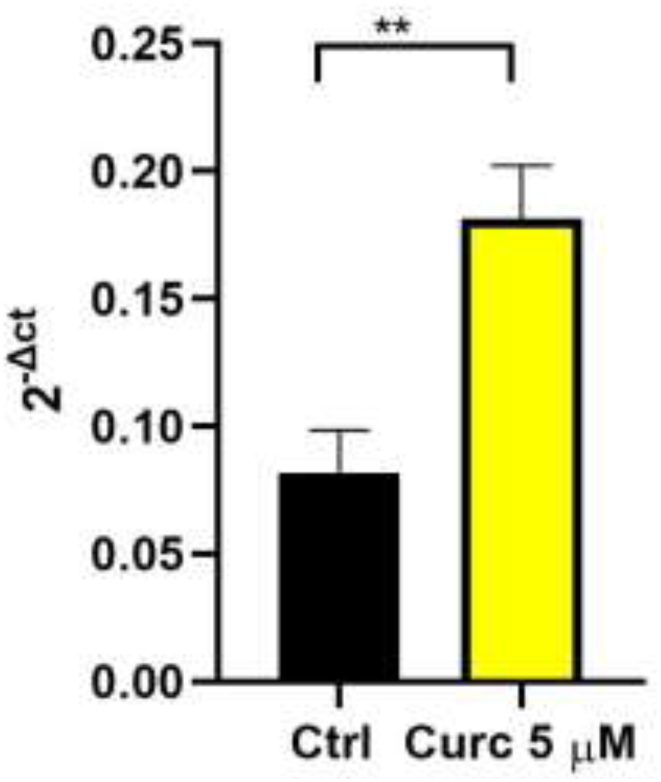
qRT PCR in differentiated HL-60 cells treated with Curcumin. After differentiation, HL-60 cells were incubated with 0.1% DMSO (Ctrl) or 5 µM Curc for 2 h. RNA was extracted and reverse transcribed, and the specific sequences for the HO transcript were amplified and quantified by RT-PCR. The bars indicate the average of three quadruplicate experiments with values expressed as 2^-ΔCt^ ± s.d. Symbols indicate significance calculated with T-Test Students: Symbols ** indicate p<0.01.

**Table S1.**
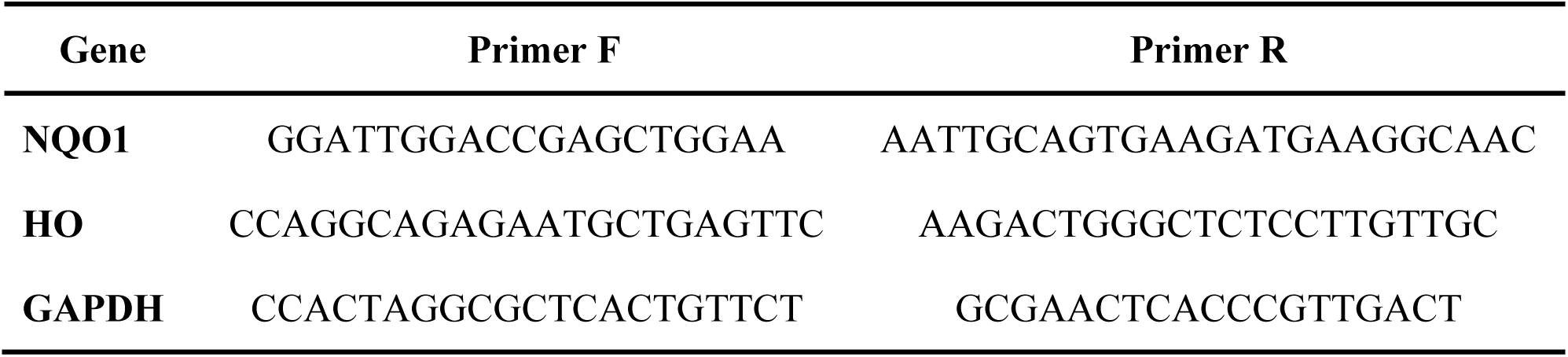
qRT-PCR primers.

## References

Calabrese, E.J., Bachmann, K.A., Bailer, A.J., Bolger, P.M., Borak, J., Cai, L., et al., 2007. Biological stress response terminology: Integrating the concepts of adaptive response and preconditioning stress within a hormetic dose-response framework. Toxicol Appl Pharmacol 222, 122–8. https://10.1016/j.taap.2007.02.015

Calabrese, E.J., Osakabe, N., Di Paola, R., Siracusa, R., Fusco, R., D’Amico, R., et al., 2023. Hormesis defines the limits of lifespan. Ageing Res Rev 91, 102074. https://10.1016/j.arr.2023.102074

Carvalho, H., 2019. Air pollution-related deaths in Europe - time for action. J Glob Health 9, 020308. https://10.7189/jogh.09.020308

Cheng, A.L., Hsu, C.H., Lin, J.K., Hsu, M.M., Ho, Y.F., Shen, T.S., et al., 2001. Phase I clinical trial of curcumin, a chemopreventive agent, in patients with high-risk or pre-malignant lesions. Anticancer Res 21, 2895–900.

Chun, K.S., Raut, P.K., Kim, D.H., Surh, Y.J., 2021. Role of chemopreventive phytochemicals in NRF2-mediated redox homeostasis in humans. Free Radic Biol Med 172, 699–715. https://10.1016/j.freeradbiomed.2021.06.031

Daniel, S., Limson, J.L., Dairam, A., Watkins, G.M., Daya, S., 2004. Through metal binding, curcumin protects against lead- and cadmium-induced lipid peroxidation in rat brain homogenates and against lead-induced tissue damage in rat brain. J Inorg Biochem 98, 266–75. https://10.1016/j.jinorgbio.2003.10.014

Della Ragione, F., Cucciolla, V., Criniti, V., Indaco, S., Borriello, A., Zappia, V., 2003. p21Cip1 gene expression is modulated by Egr1: a novel regulatory mechanism involved in the resveratrol antiproliferative effect. J Biol Chem 278, 23360–8. https://10.1074/jbc.M300771200

Ebrahimi, M., Khalili, N., Razi, S., Keshavarz-Fathi, M., Rezaei, N., 2020. Effects of lead and cadmium on the immune system and cancer progression. J Environ Health Sci Eng 18, 335–343. https://10.1007/s40201-020-00455-2

Esposito, F., Nardone, A., Fasano, E., Scognamiglio, G., Esposito, D., Agrelli, D., et al., 2018. A systematic risk characterization related to the dietary exposure of the population to potentially toxic elements through the ingestion of fruit and vegetables from a potentially contaminated area. A case study: The issue of the “Land of Fires” area in Campania region, Italy. Environ Pollut 243, 1781–1790. https://10.1016/j.envpol.2018.09.058

Eybl, V., Kotyzova, D., Koutensky, J., 2006. Comparative study of natural antioxidants - curcumin, resveratrol and melatonin - in cadmium-induced oxidative damage in mice. Toxicology 225, 150–6. https://10.1016/j.tox.2006.05.011

Gallagher, R., Collins, S., Trujillo, J., McCredie, K., Ahearn, M., Tsai, S., et al., 1979. Characterization of the continuous, differentiating myeloid cell line (HL-60) from a patient with acute promyelocytic leukemia. Blood 54, 713–33.

He, W.J., Lv, C.H., Chen, Z., Shi, M., Zeng, C.X., Hou, D.X., et al., 2023. The Regulatory Effect of Phytochemicals on Chronic Diseases by Targeting Nrf2-ARE Signaling Pathway. Antioxidants (Basel) 12. https://10.3390/antiox12020236

Iacomino, G., Tecce, M.F., Grimaldi, C., Tosto, M., Russo, G.L., 2001. Transcriptional response of a human colon adenocarcinoma cell line to sodium butyrate. Biochem Biophys Res Commun 285, 1280–9. https://10.1006/bbrc.2001.5323

Jacob, S., Nair, A.B., Shah, J., Gupta, S., Boddu, S.H.S., Sreeharsha, N., et al., 2022. Lipid Nanoparticles as a Promising Drug Delivery Carrier for Topical Ocular Therapy-An Overview on Recent Advances. Pharmaceutics 14. https://10.3390/pharmaceutics14030533

James, S.Y., Williams, M.A., Kelsey, S.M., Newland, A.C., Colston, K.W., 1997. The role of vitamin D derivatives and retinoids in the differentiation of human leukaemia cells. Biochem Pharmacol 54, 625–34. https://10.1016/s0006-2952(97)00195-0

Komatsu, M., Kurokawa, H., Waguri, S., Taguchi, K., Kobayashi, A., Ichimura, Y., et al., 2010. The selective autophagy substrate p62 activates the stress responsive transcription factor Nrf2 through inactivation of Keap1. Nat Cell Biol 12, 213–23. https://10.1038/ncb2021

Lee, B., Kim, Y.H., Lee, W., Choi, H.Y., Lee, J., Kim, J., et al., 2023. USP13 deubiquitinates p62/SQSTM1 to induce autophagy and Nrf2 release for activating antioxidant response genes. Free Radic Biol Med 208, 820–832. https://10.1016/j.freeradbiomed.2023.09.024

Lee, H.W., Choi, J.H., Seo, D., Gavaachimed, L., Choi, J., Park, S., et al., 2024. EGCG-induced selective death of cancer cells through autophagy-dependent regulation of the p62-mediated antioxidant survival pathway. Biochim Biophys Acta Mol Cell Res 1871, 119659. https://10.1016/j.bbamcr.2024.119659

Lin, X., Bai, D., Wei, Z., Zhang, Y., Huang, Y., Deng, H., et al., 2019. Curcumin attenuates oxidative stress in RAW264.7 cells by increasing the activity of antioxidant enzymes and activating the Nrf2-Keap1 pathway. PLoS One 14, e0216711. https://10.1371/journal.pone.0216711

Liu, B.Q., Gao, Y.Y., Niu, X.F., Xie, J.S., Meng, X., Guan, Y., et al., 2010. Implication of unfolded protein response in resveratrol-induced inhibition of K562 cell proliferation. Biochem Biophys Res Commun 391, 778–82. https://10.1016/j.bbrc.2009.11.137

Liu, W., Xu, Z., Li, H., Guo, M., Yang, T., Feng, S., et al., 2017. Protective effects of curcumin against mercury-induced hepatic injuries in rats, involvement of oxidative stress antagonism, and Nrf2-ARE pathway activation. Hum Exp Toxicol 36, 949–966. https://10.1177/0960327116677355

Livak, K.J., Schmittgen, T.D., 2001. Analysis of relative gene expression data using real-time quantitative PCR and the 2(-Delta Delta C(T)) Method. Methods 25, 402–8. https://10.1006/meth.2001.1262

Lozzio, B.B., Lozzio, C.B., 1979. Properties and usefulness of the original K-562 human myelogenous leukemia cell line. Leuk Res 3, 363–70. https://10.1016/0145-2126(79)90033-x

Miyaura, C., Abe, E., Suda, T., Kuroki, T., 1985. Alternative differentiation of human promyelocytic leukemia cells (HL-60) induced selectively by retinoic acid and 1 alpha,25-dihydroxyvitamin D3. Cancer Res 45, 4244–8.

Mizushima, N., 2020. The ATG conjugation systems in autophagy. Curr Opin Cell Biol 63, 1–10. https://10.1016/j.ceb.2019.12.001

Mizushima, N., Levine, B., 2020. Autophagy in Human Diseases. The New England journal of medicine 383, 1564–1576. https://10.1056/NEJMra2022774

Mohajeri, M., Rezaee, M., Sahebkar, A., 2017. Cadmium-induced toxicity is rescued by curcumin: A review. Biofactors 43, 645–661. https://10.1002/biof.1376

Moosavi, M.A., Haghi, A., Rahmati, M., Taniguchi, H., Mocan, A., Echeverría, J., et al., 2018. Phytochemicals as potent modulators of autophagy for cancer therapy. Cancer Lett 424, 46–69. https://10.1016/j.canlet.2018.02.030

Musial, C., Siedlecka-Kroplewska, K., Kmiec, Z., Gorska-Ponikowska, M., 2021. Modulation of Autophagy in Cancer Cells by Dietary Polyphenols. Antioxidants (Basel) 10. https://10.3390/antiox10010123

Nelson, K.M., Dahlin, J.L., Bisson, J., Graham, J., Pauli, G.F., Walters, M.A., 2017. The Essential Medicinal Chemistry of Curcumin. J Med Chem 60, 1620–1637. https://10.1021/acs.jmedchem.6b00975

Nimiya, Y., Wang, W., Du, Z., Sukamtoh, E., Zhu, J., Decker, E., et al., 2016. Redox modulation of curcumin stability: Redox active antioxidants increase chemical stability of curcumin. Mol Nutr Food Res 60, 487–94. https://10.1002/mnfr.201500681

Peterson, G.L., 1979. Review of the Folin phenol protein quantitation method of Lowry, Rosebrough, Farr and Randall. Anal Biochem 100, 201–20. https://10.1016/0003-2697(79)90222-7

Rana, M.N., Tangpong, J., Rahman, M.M., 2018. Toxicodynamics of Lead, Cadmium, Mercury and Arsenic-induced kidney toxicity and treatment strategy: A mini review. Toxicol Rep 5, 704–713. https://10.1016/j.toxrep.2018.05.012

Rodrigue, C.M., Arous, N., Bachir, D., Smith-Ravin, J., Romeo, P.H., Galacteros, F., et al., 2001. Resveratrol, a natural dietary phytoalexin, possesses similar properties to hydroxyurea towards erythroid differentiation. Br J Haematol 113, 500–7. https://10.1046/j.1365-2141.2001.02746.x

Russo, G.L., Spagnuolo, C., Russo, M., 2024. Reassessing the role of phytochemicals in cancer chemoprevention. Biochem Pharmacol 116165. https://10.1016/j.bcp.2024.116165

Russo, G.L., Tedesco, I., Spagnuolo, C., Russo, M., 2017. Antioxidant polyphenols in cancer treatment: Friend, foe or foil? Semin Cancer Biol 46, 1–13. https://10.1016/j.semcancer.2017.05.005

Russo, M., Milito, A., Spagnuolo, C., Carbone, V., Rosen, A., Minasi, P., et al., 2017. CK2 and PI3K are direct molecular targets of quercetin in chronic lymphocytic leukaemia. Oncotarget 8, 42571–42587. https://10.18632/oncotarget.17246

Russo, M., Moccia, S., Luongo, D., Russo, G.L., 2023. Senolytic Flavonoids Enhance Type-I and Type-II Cell Death in Human Radioresistant Colon Cancer Cells through AMPK/MAPK Pathway. Cancers (Basel) 15. https://10.3390/cancers15092660

Russo, M., Moccia, S., Spagnuolo, C., Tedesco, I., Russo, G.L., 2022. Carotenoid-Enriched Nanoemulsions and γ-Rays Synergistically Induce Cell Death in a Novel Radioresistant Osteosarcoma Cell Line. Int J Mol Sci 23. https://10.3390/ijms232415959

Russo, M., Nigro, P., Rosiello, R., D’Arienzo, R., Russo, G.L., 2007. Quercetin enhances CD95- and TRAIL-induced apoptosis in leukemia cell lines. Leukemia 21, 1130–3. https://10.1038/sj.leu.2404610

Russo, M., Russo, G.L., 2018. Autophagy inducers in cancer. Biochem Pharmacol 153, 51–61. https://10.1016/j.bcp.2018.02.007

Russo, M., Spagnuolo, C., Moccia, S., Tedesco, I., Lauria, F., Russo, G.L., 2021. Biochemical and Cellular Characterization of New Radio-Resistant Cell Lines Reveals a Role of Natural Flavonoids to Bypass Senescence. Int J Mol Sci 23. https://10.3390/ijms23010301

Shin, J.W., Chun, K.S., Kim, D.H., Kim, S.J., Kim, S.H., Cho, N.C., et al., 2020. Curcumin induces stabilization of Nrf2 protein through Keap1 cysteine modification. Biochem Pharmacol 173, 113820. https://10.1016/j.bcp.2020.113820

Shome, S., Talukdar, A.D., Choudhury, M.D., Bhattacharya, M.K., Upadhyaya, H., 2016. Curcumin as potential therapeutic natural product: a nanobiotechnological perspective. J Pharm Pharmacol 68, 1481–1500. https://10.1111/jphp.12611

Tang, L., Lan, J., Jiang, X., Huang, R., Pang, Q., Wu, S., et al., 2022. Curcumin antagonizes inflammation and autophagy induced by arsenic trioxide through immune protection in duck spleen. Environ Sci Pollut Res Int 29, 75344–75355. https://10.1007/s11356-022-20691-3

Triassi, M., Alfano, R., Illario, M., Nardone, A., Caporale, O., Montuori, P., 2015. Environmental pollution from illegal waste disposal and health effects: a review on the “triangle of death”. Int J Environ Res Public Health 12, 1216–36. https://10.3390/ijerph120201216

Wahyudi, L.D., Yu, S.H., Cho, M.K., 2022. The effect of curcumin on the cadmium-induced mitochondrial apoptosis pathway by metallothionein 2A regulation. Life Sci 310, 121076. https://10.1016/j.lfs.2022.121076

Wei, L., Li, S., Ma, Y., Ye, S., Yuan, Y., Zeng, Y., et al., 2024. Curcumin attenuates diphenyl phosphate-induced apoptosis in GC-2spd(ts) cells through activated autophagy via the Nrf2/P53 pathway. Environ Toxicol 39, 2032–2042. https://10.1002/tox.24092

White, E., Mehnert, J.M., Chan, C.S., 2015. Autophagy, Metabolism, and Cancer. Clin Cancer Res 21, 5037–46. https://10.1158/1078-0432.CCR-15-0490

Xu, G., Chen, H., Cong, Z., Wang, R., Li, X., Xie, Y., et al., 2024. Promotion of transcription factor EB-dependent autophagic process by curcumin alleviates arsenic-caused lung oxidative stress and inflammation in mice. J Nutr Biochem 125, 109550. https://10.1016/j.jnutbio.2023.109550

Zaccaroni, A., Corteggio, A., Altamura, G., Silvi, M., Di Vaia, R., Formigaro, C., et al., 2014. Elements levels in dogs from “triangle of death” and different areas of Campania region (Italy). Chemosphere 108, 62–9. https://10.1016/j.chemosphere.2014.03.041

Zhai, Q., Narbad, A., Chen, W., 2015. Dietary strategies for the treatment of cadmium and lead toxicity. Nutrients 7, 552–71. https://10.3390/nu7010552

Zhong, Z., Umemura, A., Sanchez-Lopez, E., Liang, S., Shalapour, S., Wong, J., et al., 2016. NF-κB Restricts Inflammasome Activation via Elimination of Damaged Mitochondria. Cell 164, 896–910. https://10.1016/j.cell.2015.12.057

